# *Trypanosoma brucei*: A novel system for metacyclogenesis and complete cyclical development *in vitro*

**DOI:** 10.1101/2022.01.18.476722

**Authors:** Beatrice Nickel, Monika Fasler, Reto Brun, Ronald Kaminsky, Isabel Roditi, Leo Jenni

## Abstract

We describe an *in vitro* system that allows metacyclogenesis of *T. brucei* ssp. and is not restricted to certain parasite lines. Following the work of Hirumi and co-workers we developed a modified *in vitro* system which allows metacyclogenesis within 7 to 20 days. The most striking changes were the addition of citrate and cis-aconitate, which triggers transformation from bloodstream to procyclic forms, and the elimination of the remaining bloodstream forms by salicylhydroxamic acid and glycerol. A positive feature of this system is that *T. b. gambiense* samples isolated from patients, and transformed to procyclic forms for transport, could give rise to infective forms that could be passaged in rodents.

## Introduction

The complex life cycle of sleeping sickness trypanosomes, which belong to the species *Trypanosoma brucei* is divided between the vertebrate and the tsetse fly host. During the course of their life cycle, these parasites encounter many differing environments and respond to these by changes in cell shape, metabolism and pattern of gene expression. Many of these life cycle transitions can be carried out *in vitro*, allowing their underlying controls to be studied.

There have been significant advances in the understanding of the control of differentiation in bloodstream forms due to the ability to culture these forms. Furthermore, studies of the sequence of events during the transformation of bloodstream to procyclic forms have precisely mapped the steps of cell differentiation (Szöőr et al., 2020). This process is supported by a special culture system using a semi-defined medium and tricarboxylic acid cycle intermediates, especially cis-aconitate (Brun and Schönenberger, 1981).

The less studied part of the life cycle, which occurs in the tsetse fly, spans the transition from non-infective procyclic forms in the midgut to epimastigote forms in the salivary glands and finally to metacyclic forms which are infectious for vertebrates. During this process, which is known as metacyclogenesis, a series of complex changes occur. These include morphological and ultrastructural modifications, activation and repression of major metabolic pathways, alterations of endocytic processes and cell cycle arrest in the metacyclic form (Natesan et al., 2007; Sharma et al., 2009; Toh et al., 2021; Vickerman, 1965).

For several species of the genus *Trypanosoma*, including *Trypanosoma brucei*, *in vitro* culture systems have been developed which allow metacyclogenesis to occur to some extent. The development of metacyclic forms of *T. b. brucei*, in culture systems which included head and salivary glands from tsetse flies, was first described by Cunningham and Honigberg (Cunningham, 1977; Cunningham and Honigberg, 1977). The yield of vertebrate infective forms produced in this system and modified versions was very low (0.01-3% of the total trypanosome population), but the infective trypanosomes produced appeared to be metacyclic forms. This culture system, although laborious in requiring organs and tissue from tsetse flies, was the first culture system in which metacyclic forms of *T. b. brucei* produced *in vitro* could be separated from non-infective forms. Subsequently, Cunningham and Kaminsky developed a culture system in which tsetse explants were replaced by a cell line from *Anopheles gambiae* (Kaminsky et al., 1987). When procyclic forms of three stocks of *T. b. brucei* and one stock of *T. b. rhodesiense* were grown at 27°C, some of them developed into forms infective for mice, and 2.8×10^5^ metacyclic forms per ml of culture supernatant could be harvested. Another culture system, which entails several steps of sub-cultivation and system changes, was described by Hirumi et al. 1992 (Hirumi et al., 1992). In this case *T. b. brucei* metacyclic forms developed in the presence of mammalian feeder layer cells and Cytodex 3 beads. This study indicates that the presence of insect tissues and cells is not necessary for metacyclogenesis. However, the yield of metacyclic forms was much lower than in the other systems and yielded approximately 5×10^4^ metacyclic forms per ml of culture per day. These *in vitro* culture systems for *T. brucei* have laid the groundwork for metacyclogenesis *in vitro*, although it was not reproducible (single events) and occurred only with selected parasite isolates and stocks. Although it is now possible to induce metacyclogenesis by ectopic expression of the RNA-binding protein RBP6, this requires genetic manipulation before a cell line can be used, and the metacyclic forms generated in this way cannot switch to bloodstream forms *in vitro* (Kolev et al., 2012; Shi et al., 2018; Toh et al., 2021).

A culture system which allows reproducible metacyclogenesis for a broad range of trypanosome isolates and stocks would eliminate the laborious work with tsetse flies and would allow the complete life cycle of *T. brucei* isolates to be studied *in vitro*. Furthermore, such a system would be a valuable tool to study life cycle transitions, e.g. genetic recombination *in vitro*, as genetic recombination, expression of meiotic markers and gamete-like forms occur in the fly salivary glands (Gibson, 2015; Gibson and Whittington, 1993; Jenni et al., 1986; Peacock et al., 2014). Here we describe a culture system for *in vitro* metacyclogenesis of the complete life cycle of *T. brucei* without the need of tsetse fly tissue.

## Materials and methods

### *Trypanosoma brucei* stocks

#### T. brucei brucei

Clone STIB 247 LA is a derivative of the primary isolate STIB 247, isolated from Coke’s hartebeest in the Serengeti National Park, Tanzania, in 1971 (Jenni et al. 1986). STIB 829 is an isolate from a tsetse fly in Uganda, in 1990 (Degen et al., 1995; Gibson, 2015).

#### *T. brucei gambiense* type 2

Clone STIB 386 AAA is derived from TH114/78 E (020), isolated from a patient in the Ivory Coast in 1978 (Jenni et al. 1986).

#### *T. brucei gambiense* type 1

STIB 887, STIB 889 and STIB 890 are stocks of procyclic forms obtained from bloodstream forms originally isolated form patients in north-western Uganda (Balmer and Caccone, 2008).

#### T. brucei rhodesiense

STIB 798 is a derivative of the primary isolate KETRI 2694 (Kagira and Maina, 2007).

### Feeder layer cells and media

Commercially available rat skeletal muscle myoblast (L6) cells (ATCC^®^ CRL-1458), human foetal lung fibroblasts (WI-38, ATCC^®^ CCL-75) and mouse embryonic fibroblast (MEF) cells (ATCC^®^ CRL-2214) were used as feeder layer cells. Feeder layers were grown in MEM medium supplemented with 15% heat inactivated foetal bovine serum (FBS). Cultures were kept at 37°C in a humidified atmosphere with 5% CO_2_.

Procyclic forms were grown in SM medium supplemented with 10% heat inactivated FBS (Cunningham, 1977). Bloodstream forms were grown in BMEM (Baltz et al., 1985) supplemented with 15% heat inactivated horse serum.

### *In vitro* transformation of bloodstream to procyclic forms

Mice were infected by intraperitoneal injection with bloodstream forms. At the first peak of parasitaemia blood was taken by heart puncture, diluted 1:2 in phosphate-buffered saline-glucose (PSG 6:4, 1% glucose) with 100iU/ml Heparin and trypanosomes were isolated by differential centrifugation at 70xg for 15 min. Supernatant with trypanosomes was centrifuged at 1250xg for 10 min. The pellet was resuspended in SM medium supplemented with 15% heat inactivated FBS and differentiation to procyclic forms was initiated by addition of 3mM citrate and 3 mM cis-aconitate (CCA) according to Brun and Schönenberger (Brun and Schönenberger, 1981). Cultures were maintained at 27°C for 7 days and then treated with 10mM salicylhydroxamic acid (SHAM) and 25mM glycerol for three hours to eliminate residual bloodstream form trypanosomes (Clarkson and Brohn, 1976). In order to check whether vertebrate infective forms survived the treatment, 10^6^ parasites were injected into CD1 mice (*T. b. brucei* or *T. b. rhodesiense*) or *Mastomys coucha* (*T. b. gambiense*).

### *In vitro* production of metacyclic forms

Procyclic forms (PCF) were added to T-12.5 flasks (10^7^ PCF/flask) containing feeder layer cells and 5ml MEM medium without glucose, but supplemented with 10mM proline, 2mM glutamine and 15% heat inactivated FBS. The co-cultures were kept at 27°C in a humidified atmosphere with 5% CO_2_. After 3 days the medium containing all the non-attached trypanosomes was discarded and replaced by 5ml fresh medium.

Seven to seventeen days later, depending on the trypanosome stock, the first metacyclic forms could be separated from non-infective culture forms by means of DEAE-cellulose (Whatman Chromedia DE 52) column chromatography according to Gardiner et al. (Gardiner et al., 1980). One aliquot of the eluted trypanosomes was then added to 300μl of 0.9% NaCl solution and injected into CD1 mice to test the infectivity and the other aliquot was added to 200μl of BMEM medium to propagate bloodstream forms *in vitro*.

## Results

The culture system was initially established using STIB 386 and STIB 247. Depending on the trypanosome stock infective metacyclic forms appeared between day 8 and day 20 of co-cultivation with the L6 feeder layer cells (Table 1). To test infectivity, trypanosomes eluted from a DEAE cellulose column were injected into CD1 mice (*Trypanosoma b. brucei*) or *Mastomys coucha* (*T. b. gambiense*). Parasites were found in the blood 5 to 15 days post infection.

**Table 1:** The *in vitro* culture system was tested with nine different strains. All three subspecies of *T. brucei* are represented. f.l. = feeder layer; * passaged on a second feeder layer. BSF, bloodstream form. nd, not done.

| Strain | Subspecies | First metacyclic forms | Infectivity | BSF culture |
| --- | --- | --- | --- | --- |
| STIB 386 | *T. b. gambiense* | d10 | + | + axenic |
| STIB 247 | *T. b. brucei* | d18* | + | + axenic |
| STIB 829 | *T. b. brucei* | d20* | + | nd |
| Patient derived | *T. b. gambiense* | d10 | + | - |
| STIB 887 | *T. b. gambiense* | d9 | + | 5d with f.l. |
| STIB 889 | *T. b. gambiense* | d11 | + | 4d with f.l. |
| STIB 890 | *T. b. gambiense* | d8 | + | 7d with f.l. |
| STIB 798 | *T. b. rhodesiense* | d8 | + | nd |

It was also possible to complete the life cycle completely *in vitro*. Trypanosomes purified by DEAE cellulose chromatography were taken directly into culture and maintained in MEM supplemented with Baltz stock at 37°C and 5% CO_2_. (B_a_ltz _e_t al., 1985) The three *T. b. gambiense* strains STIB 887, STIB 889 and STIB 890 were very difficult to maintain in culture directly after cellulose chromatography. Even in the presence of a feeder layer, they survived for only a few days and hardly proliferated (see Table 1). However, infective forms from strains already adapted to axenic culture conditions before going through the whole life cycle *in vitro* could be maintained as axenic cultures in BMEM.

With some of these strains, *in vitro* metacyclogenesis was performed for a second and even a third time. The time elapsing until the first infective parasites could be purified became shorter with each cycle, indicating that some form of selection of takes place (Table 3). It is also worth noting that procyclic forms maintained for more than three months in culture were still capable of completing the whole cycle *in vitro*.

**Table 2:** STIB 386 was cultivated on different feeder layers. MCF, metacyclic forms; BSF, bloodstream forms. L6, rat myoblasts; WI-38, human foetal lung fibroblasts; MEF, mouse embryonic fibroblasts.

| Feeder layer | First MCF | peak | titre | infectivity | BSF culture |
| --- | --- | --- | --- | --- | --- |
| L6 | d10 | d13 | 2x10<sup>5</sup>/ml | + | + axenic |
| WI-38 | d13 | nd | 100/ml | + | - |
| MEF | d7 | d9 | 1x10<sup>3</sup>/ml | + | + axenic |

**Table 3:** Appearance of metacyclic forms of STIB 386 during three cycles on L6 feeder layers. MCF, metacyclic forms.

| Cycle | First MCF | Peak |
| --- | --- | --- |
| 1 | d10 | d13 |
| 2 | d7 | d10 |
| 3 | d5 | d8 |

To keep the trypanosomes under optimal conditions, they were transferred to a new feeder layer every 10 to 15 days depending on the state of the mammalian cells. With each change, only non-attached parasites were transferred to the new feeder layer which explains a period of one to two days in which no new infective parasites could be detected. Furthermore, the number of parasites eluted from the DEAE cellulose column decreases the longer the culture is maintained (see Table 4). The most metacyclic forms can be observed 2 to 5 days after the first time they appear. For some strains, which can take up to three weeks to develop infective parasites, this may entail maintaining them on sequential feeder layers.

**Table 4:** STIB 386 on L6 feeder layer, 3 passages to fresh L6 feeder cells, one cycle. MCF, metacyclic forms.

| Feeder layer | First MCF | Titre |
| --- | --- | --- |
| 1st | d10 | $2 \times 10^5/\text{ml}$ |
| 2nd | d2 | $1 \times 10^4/\text{ml}$ |
| 3rd | d3 | $0.5 \times 10^3/\text{ml}$ |

